# Natural variation in the regulation of neurodevelopmental genes modifies flight performance in *Drosophila*

**DOI:** 10.1101/2020.05.27.118604

**Authors:** Adam N. Spierer, Jim A. Mossman, Samuel Pattillo Smith, Lorin Crawford, Sohini Ramachandran, David M. Rand

## Abstract

The winged insects of the order *Diptera* are colloquially named for their most recognizable phenotype: flight. These insects rely on flight for a number of important life history traits, like dispersal, foraging, and courtship. Despite the importance of flight, relatively little is known about the genetic architecture of variation for flight performance. Accordingly, we sought to uncover the genetic modifiers of flight using a measure of flies’ reaction and response to an abrupt drop in a vertical flight column. We conducted an association study using 197 of the *Drosophila* Genetic Reference Panel (DGRP) lines, and identified a combination of additive and marginal variants, epistatic interactions, whole genes, and enrichment across interaction networks. We functionally validated 13 of these candidate genes’ (*Adgf-A/Adgf-A2/CG32181, bru1, CadN, CG11073, CG15236, CG9766, CREG, Dscam4, form3, fry, Lasp/CG9692, Pde6, Snoo*) contribution to flight, two of which (*fry* and *Snoo*) also validate a whole gene analysis we introduce for the DGRP: `PEGASUS_flies`. Overall, our results suggest modifiers of muscle and wing morphology, and peripheral and central nervous system assembly and function are all important for flight performance. Additionally, we identified *ppk23*, an Acid Sensing Ion Channel (ASIC) homolog, as an important hub for epistatic interactions. These results represent a snapshot of the genetic modifiers affecting drop-response flight performance in *Drosophila*, with implications for other insects. It also draws connections between genetic modifiers of performance and BMP signaling and ASICs as targets for treating neurodegeneration and neurodysfunction.

**Author summary:** Insect flight is a widely recognizable phenotype of winged insects, hence the name: flies. While fruit flies, or *Drosophila melanogaster*, are a genetically tractable model, flight performance is a highly integrative phenotype, making it challenging to comprehensively identify the genetic modifiers that contribute to its genetic architecture. Accordingly, we screened 197 Drosophila Genetic Reference Panel lines for their ability to react and respond to an abrupt drop. Using several computational tools, we successfully identified several additive, marginal, and epistatic variants, as well as whole genes and altered sub-networks of gene-gene and protein-protein interaction networks, demonstrating the benefits of using multiple methodologies to elucidate the genetic architecture of complex traits more generally. Many of these significant genes and variants mapped to regions of the genome that affect development of sensory and motor neurons, wing and muscle development, and regulation of transcription factors. We also introduce PEGASUS_flies, a *Drosophila*-adapted version of the PEGASUS platform first used in human studies, to infer gene-level significance of association based on the distribution of individual variant *P*-values. Our results contribute to the debate over the relative importance of individual, additive factors and epistatic, or higher order, interactions, in the mapping of genotype to phenotype.

## Introduction

Flight is one of the most distinguishing features of winged insects, especially the taxonomic order *Diptera*. Colloquially named “flies,” these insects rely on their namesake for many facets of their life history: dispersal, foraging, evasion, migration, and mate finding [1]. Because flight is central to flies’ life history, many of the most critical genes for flight are strongly conserved [2, 3].

These “flight-critical” genes are necessary for flight, even as the structures and neural circuits they form are co-opted for other phenotypes, like courtship song and display [4, 5]. For example, *Wingless* is an important developmental patterning gene necessary for wing formation [6] and *Act88F* is one of the main actin isoforms in the indirect flight muscles [7]. We will designate these types of genes that play outsized roles in enabling flight “flight critical” genes, since altering their sequence or expression profile is more likely to result in large flight performance deficits. On the other hand, we will designate “flight-important” genes as those with more modest effects on flight, since they are important but not critical. In an evolutionary context, purifying selection would act on flight-critical genes more strongly than flight-important genes, meaning flight-important genes can harbor natural variants that might otherwise vary the flight phenotype. These genes are found across systems, including metabolism [8], muscle function [9], neuronal function [10, 11], and anatomical development [12, 13]. Genes filling multiple roles across systems are pleiotropic, and those with sufficient natural variation are likely to contribute to complex traits and disease [14, 15]. These traits and diseases’ independent, yet interconnected, genetic architecture make them inherently challenging to study because they are comprised of several modifiers of small to moderate effect size [16–18].

We can leverage natural variants in flight-important genes to uncover novel associations between genotype and phenotype that otherwise modify flight-critical genes’ function, via Genome Wide Association Study (GWAS). The *Drosophila* Genetics Reference Panel [19, 20] (DGRP) is a common resource for performing this type of analysis. The DGRP is a panel of 205 genetically distinct *D. melanogaster* lines represents a snapshot of natural variation. Previous studies on complex and highly polygenic, quantitative traits identify several candidate loci contributing to insect- and *Drosophila*-specific traits [21–23], as well as traits affecting human health and disease [24–27].

Accordingly, this study was designed to identify the genetic modifiers of flight performance and map the underlying genetic architecture. We screened males and females from 197 of the 205 DGRP lines and analyzed both sexes, as well as the average and difference between sexes. Traditional association studies focus on the contribution of additive and dominant variants, however, these fail to identify different types of modifiers with different effect sizes. Accordingly, we took several approaches to identify modifiers at the individual variant, whole gene, and network levels. Accordingly, we identified 180 additive variants, 70 marginal variants, 12161 unique epistatic interactions, and nine interaction sub-networks containing 539 genes contributing to flight performance. We also identified 72 whole genes using `PEGASUS_flies`, a novel modification of the human-based `PEGASUS` program [28] that we modified to work with *Drosophila* and DGRP studies <https://github.com/ramachandran-lab/PEGASUS_flies> (File S4).

Taken together, our results strongly suggest variation in flight performance across natural populations is affected by cis- and trans-regulatory elements’ role in modifying 1) development of wing morphology, indirect flight musculature, and sensory organs; and the connectivity between the peripheral and central nervous systems. These results are further supported by functional validations of 13 candidate genes, many with known roles in altering neurogenesis and development. Overall, our results suggest important roles for modifiers of BMP signaling in neurodevelopment and pickpocket 23 (ppk23)—a degenerin/epithelial sodium channels (DEG/ENaC) homologous in humans with Acid Sensing Ion Channels (ASIC)—in altering affecting flight performance. These findings address an underexplored body of literature [29–32] calling for genetic and pharmacological targets of BMP signaling genes and ASIC for treating neuroinjury and neurodegenerative diseases in humans.

## Results

### Variation in flight performance across the DGRP

Cohorts of approximately 100 flies from 197 lines of the DGRP (S1 Table) were tested for flight performance using a flight column [33] (Fig 1A). We confirmed the repeatability of our assay by retesting 12 lines of varied ability reared 10 generations apart. We observed very strong agreement between generations (r = 0.95; S1 Fig), affirming a role for genetic, rather than environmental or experimental, variation in driving phenotypic variation. We recorded high-speed videos for a weak, intermediate, and strong genotype entering the flight column (Figs 1B-D; S1 File) and concluded this assay is best for studying the reaction and response to an abrupt drop. There was strong agreement between sex-pairs’ mean landing height for each genotype (r = 0.75; Fig 1E), suggesting the genetic architecture is mostly shared between the sexes. As expected, there was a modest degree of sexual dimorphism in performance, with males outperforming females (male: 0.80m ± 0.06 SD; female: 0.73m ± 0.07 SD; Fig 1F; S2 Table), though the broad sense heritability (*H^2^*) for each sex was nearly the same (*H^2^* = 13.5%; *H^2^* = 14.4%). In addition to males and females, we also investigated the phenotypic variation in the average (sex-average) and difference (sex-difference) between sexes (S2 Fig).

**Fig 1.**
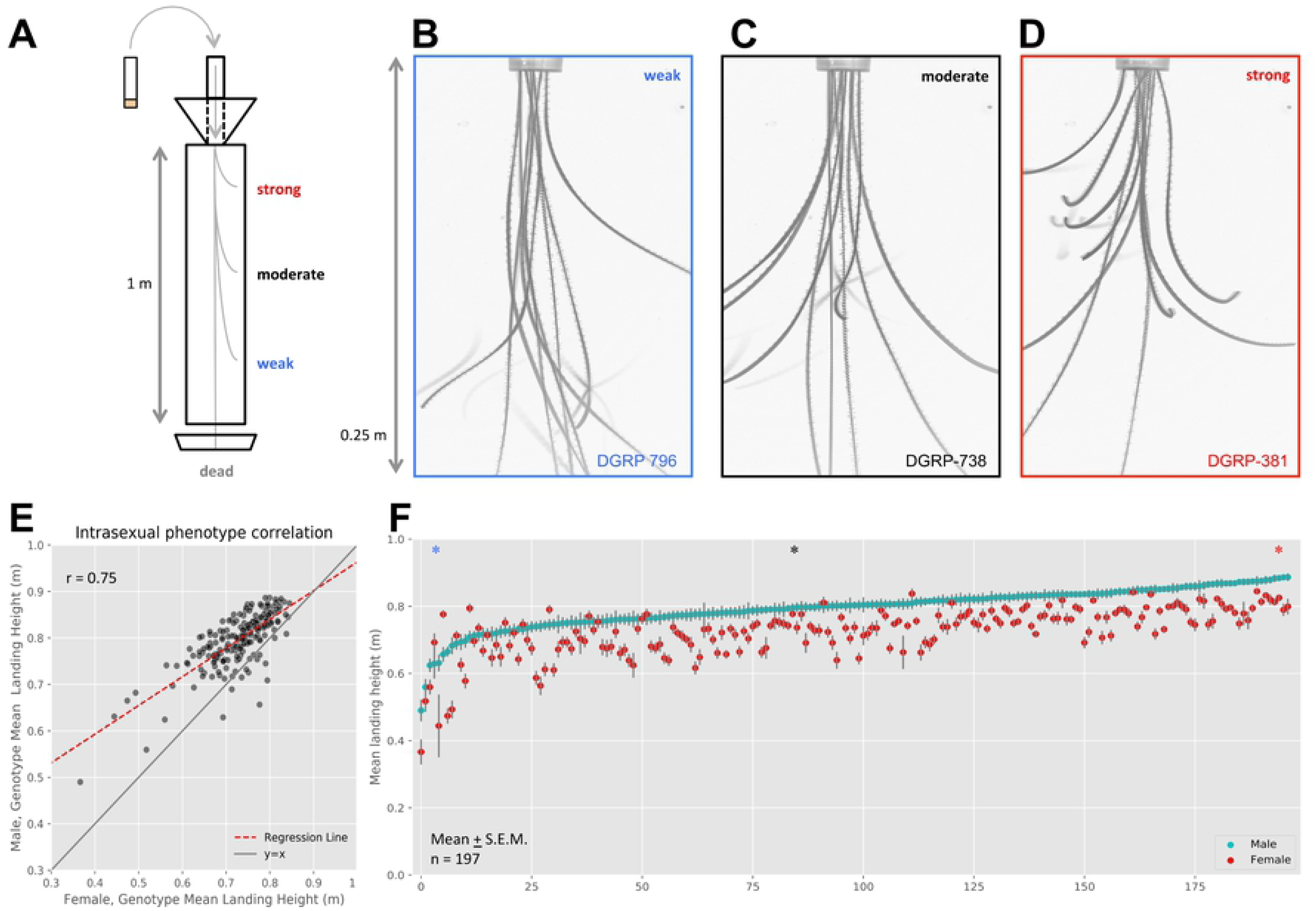
DGRP lines show differences in flight performance across lines. (A) Flight performance assay measures the average landing height of flies as they fall through a flight column. Vials of flies are sent down the top chute and abruptly stop at the bottom, ejecting flies into a meter-long column. Falling flies will instinctively right themselves and fly to the periphery, doing so at different times depending on their performance ability. (B-D) Collapsed z-stacks of high-speed video frames from the top quarter of the flight column illustrate these performance differences in (B) weak, (C) intermediate, and (D) strong genotypes. (E) There is sexual dimorphism within genotypes (deviation of red dashed regression line from y = x solid gray line), though sexes are well correlated (r = 0.75, n = 197). (F) Sexually dimorphic performances are also viewable in the distribution of performances for each male (cyan) and female (red) genotype pair (mean ± S.E.M.). Sex-genotype pairs are sorted in order of increasing male mean landing height. Genotype performances for genotypes in B-D are indicated on the distribution with the corresponding color-coded asterisk (*) above the respective genotype position.

Before running the association analysis, we tested whether flight performance was a unique phenotype. We compared our phenotype scores for males and female against publically available phenotypes on the DGRP2 webserver. We found no significant regression between flight performance and any of the phenotypes in either sex after correcting for multiple testing (*P* ≤ 1.85E-3; S3 Table). This negative result suggests our measure of flight performance is a unique phenotype among those reported.

### Association of additive SNPs with flight performance

We conducted a Genome Wide Association Study (GWAS) to identify genetic markers associated with flight performance. We performed an analysis with 1,901,174 common variants (MAF ≥ 0.05) on the additive genetic effects of four sex-based phenotypes: males, females, sex-average, and sex-difference. Some phenotypes covaried with the presence of major inversions (S4 Table), so we analyzed association results using a mixed model (Fig 2A) to account for *Wolbachia* infection status, presence of inversions, and polygenic relatedness (S3 and S4 Figs). Annotations and unreferenced descriptors of genes’ functions, expression profiles, and orthologs were gathered from autogenerated summaries on FlyBase [34, 35]. These summaries and descriptors were compiled from data supplied by the Gene Ontology Consortium [36, 37], the Berkeley *Drosophila* Genome Project [38], FlyAtlas [39], The Alliance of Genome Resources Consortium [40], modENCODE [34], PAINT[41], the DRSC Integrative Ortholog Prediction Tool (DIOPT) [42], and several transcriptomics and proteomic datasets [9, 12, 39, 43–46].

**Fig 2.**
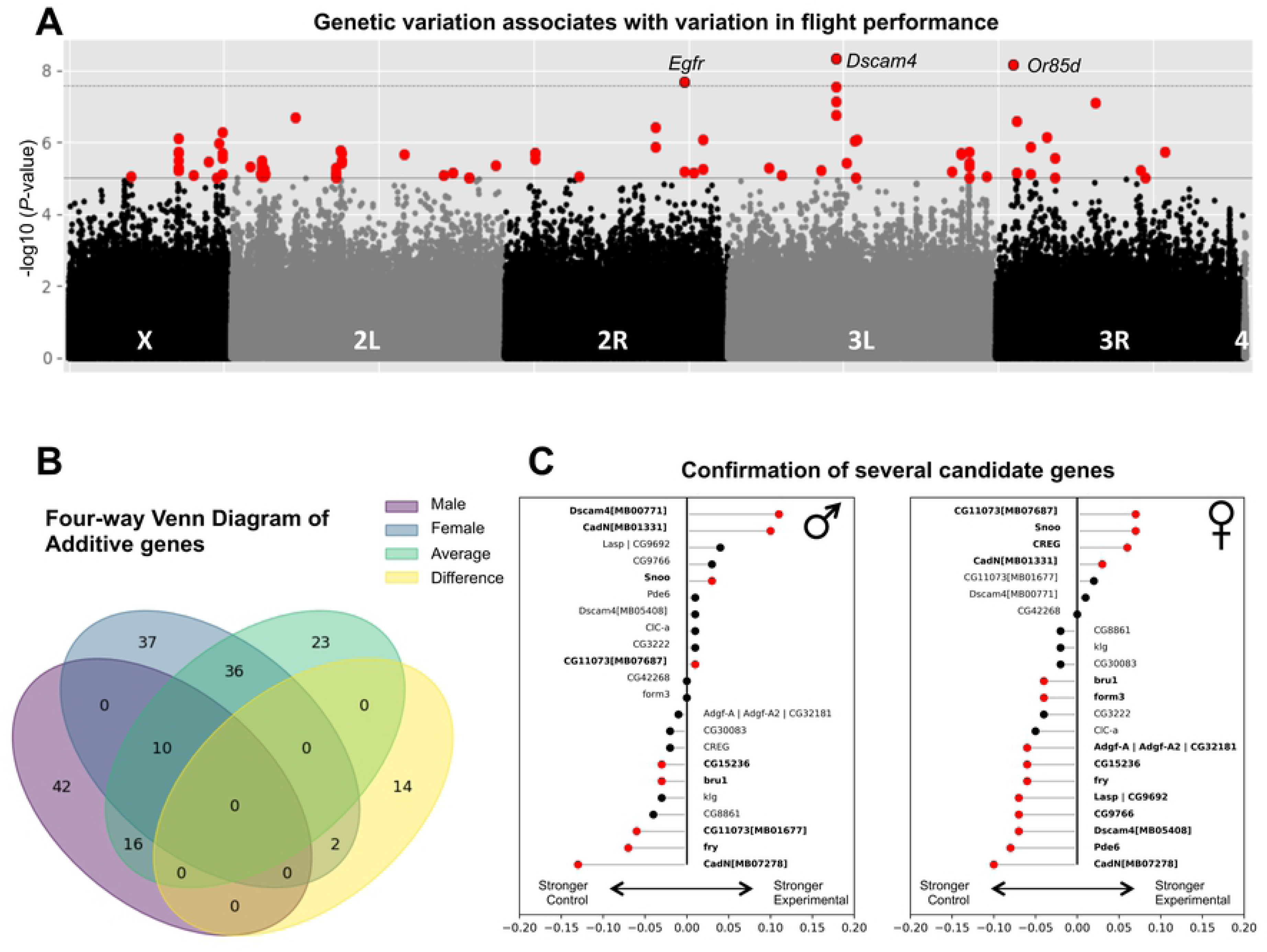
Variation in flight performance associated with several additive variants, some of which were functionally validated. (A) An additive screen for genetic variants identified several variants that exceeded the traditional DGRP (*P* ≤1E-5) threshold (gray line). These points (red points) were spread throughout the genome on all but chromosome 4. Sex-average variants pictured, though other sex-based phenotypes had similar profiles. (B) Approximately half of all variants were shared with at least one other sex-based analysis, while the other half of all variants was exclusive to a single analysis. (C) Candidate genes were selected based on the genes that the most significant variants mapped to. Both sexes were tested for flight performance. Validated genes were determined if there was a significant difference between experimental lines homozygous for an insertional mutant in the candidate gene and their background control lines lacking the insertional mutant (red points, Mann-Whitney-U test, *P* ≤ 0.05). Very significant candidate genes (*CadN*, *CG11073*/*flapper*, and *Dscam4*) each had two independent validation lines.

We filtered additive variants with a strict Bonferroni threshold (*P* ≤ 2.63E-8). Taking a MinSNP approach, which identifies significant genes if their lowest (most significant) variant *P*-value crosses a threshold [28], we identified six unique variants, five of which mapped to six genes (*CG15236*, *CG34215, Dscam4, Egfr, fd96Ca, Or85d*) (Table 1). Variants mapping to *Egfr* and *fd96Ca* also contained known embryonic cis-regulatory elements (transcription factor binding sites (TFBS) and a silencer) [47]. Of note, *Dscam4* was deemed “damaged” in 38 of the lines tested [19]; however, the difference between mean landing heights of the damaged vs. undamaged lines was less than 1 cm (*P* = 0.32, Welch’s T-test).

**Table 1.**
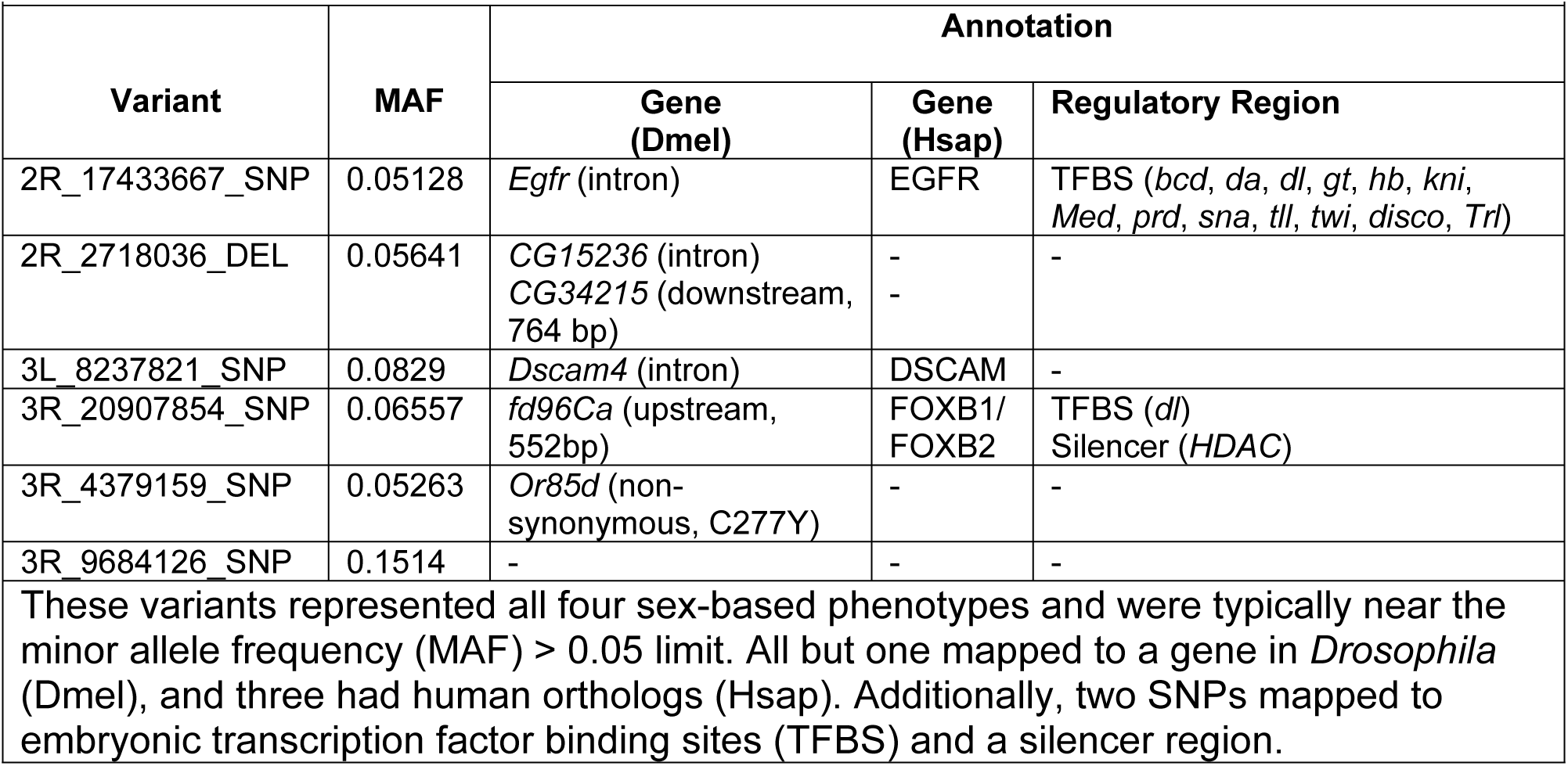
Six additive variants surpassed the Bonferroni significance threshold.

Using the traditional DGRP significance threshold (*P* ≤ 1E-5) [48], we identified 180 variants across all four sex-based phenotypes (Fig 2B, S5 Fig, S5 Table). The individual additive variant with the largest effect size contributed 0.045 meters (or 0.97% of the sum of all significant variants) for males and 0.064 meters (1.1% of the sum of all significant variants) for females. For reference, the variant with the smallest significant effect size was 1.7E-4 meters (or 0.0036% of the sum of all significant variants) for males and 5.7E-3 meters (or 0.095% of the sum of all significant variants) for females.

All but 19 variants mapped to intergenic or non-coding regions, which are generally indicative of cis-regulatory regions. Of the non-coding variants, 149 mapped to 136 unique genes across the sex-based analyses (Table 2). These included development and function of the nervous system (*aru*, *CadN*, *ChAT*, *chinmo*, *chn*, *CNMaR*, *CSN6*, *DIP-delta*, *Dscam4*, *Egfr*, *fd96Ca*, *form3*, *fry*, *hll*, *htk*, *jeb*, *kek2*, *klg*, *klu*, *Mbs*, *Mmp2*, *nompC*, *Or46a*, *Or85d*, *Pdp1*, *Ptp10D*, *pyd*, *Rbp6*, *rut*, *Sdc*, *SK*, *SKIP*, *Spn*, *Snoo*, *Tmc*), neuromuscular junction (*fend*, *Gad1*, *Gαo/Galphao*, *jeb*, *Sdc*, *Syt1*), muscle (*bru1*, *bves*, *CG17839*, *jeb*, *Lasp*, *Pdp1*, *rhea*), cuticle and wing morphogenesis (*CG15236*, *CG34220*, *CG43218*, *Egfr*, *frtz*, *fry, Tg*), endoplasmic reticulum (*CG33110*, *CG43783*, *tbc*, *Vti1b*) and Golgi body functions (*Gmap*, *Rab30*, *Vti1b*), and regulation of translation (*mip40, mxt*, *Rbm13*, *Wdr37*). Approximately half of all variants were present in two or three sex-based analyses, though the remainder were unique to one (Fig 2B). Several variants mapped to transcription factors (*Asciz*, *Camta*, *CG18011*, *chinmo*, *chn*, *Eip78C*, *fd96Ca*, *Pdp1*, *run*) broadly affecting development and neurogenesis [34, 35]. Despite the enrichment for several annotations, we failed to identify any significant gene ontology (GO) categories using GOwinda [49], a GWAS-specific gene set enrichment analysis.

**Table 2.**
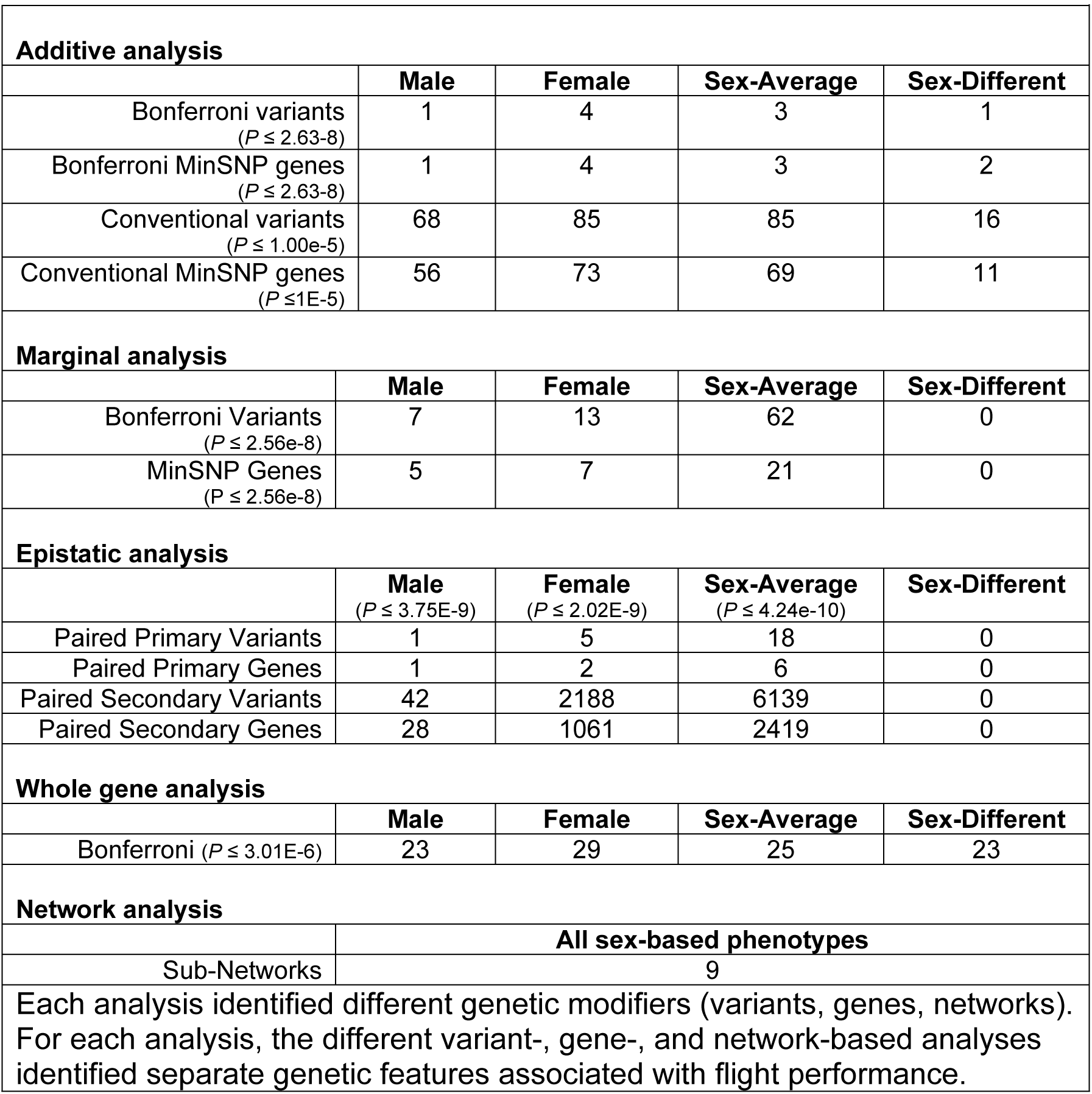
Aggregated gene and variant counts by sex-based phenotype for each analysis.

### General development and neurodevelopmental genes validated to affect flight performance

We performed functional validations on a subset of the genes mapped from variants identified in the Bonferroni and sex-average analysis. We identified 21 unique candidate genes for which a *Minos* enhancer trap *Mi{ET1}* insertional mutation line [50] was publically available [51] (S1 Table; *Adgf-A/Adgf-A2/CG32181, bru1, CadN, CG11073, CG15236, CG9766, CREG, Dscam4, form3, fry, Lasp/CG9692, Pde6, Snoo*). Three additional stocks for *CadN*, *Dscam4*, and *CG11073* were also tested for their strength of association. Finally, an insertion line for *CREG* was also included as a negative control, since it was not significant in the additive or subsequent analyses.

Candidate genes were functionally validated by comparing the distribution in mean landing heights of stocks homozygous for the insertion and their paired control counterpart (S6 Fig) using a Mann-Whitney-U test (Fig 2C; S6 Table). Several were involved in neurodevelopment (*CadN, CG9766*, *CG11073*, *CG15236, Dscam4*, *form3*, *fry*, and *Snoo*), muscle development (*bru1* and *Lasp*), and transcriptional regulation of gene expression (*Pde6* and *CREG*). Both CG9766 and CG11073 are unnamed candidate genes. In validating roles for both these genes, we are naming them *tumbler* (*tumbl*) and *flapper* (*flap*), respectively, based on the tumbling and flapping motions of weaker flies struggling to right themselves in the flight performance assay.

### Association of gene-level significance and interaction networks with flight performance

The minSNP approach on the additive variants prioritizes the identification of genes containing variants with larger effects [28]. However, this approach ignores linkage blocks and gene length, which can bias results. It is important to account for gene length because many neurodevelopmental genes can be lengthy and exceed 100kb (*CadN*, 131kb). One alternative approach is Precise, Efficient Gene Association Score Using SNPs (`PEGASUS`), which assesses whole gene significance scores based on the distribution of a gene’s variant *P*-value distributions with respect to a null chi-squared distribution [28]. This approach enriches for whole genes of moderate effect and enables the identification of genes that might go undetected in a minSNP approach.

While `PEGASUS` is configured for human populations, we developed `PEGASUS_flies`, a modified version for *Drosophila* <https://github.com/ramachandran-lab/PEGASUS_flies>. This platform is configured to work with DGRP data sets, and can be customized to accept other *Drosophila*-based or model screening panels. From our additive variants, `PEGASUS_flies` identified 72 unique genes across the all sex-based phenotypes, whose gene scores passed a Bonferroni threshold (*P* ≤ 3.03E-6; S7 Table). These genes were present on five of the six chromosome arms tested (Fig 3A). They were generally different from those identified in the additive approach’s minSNP analyses (Fig 3B, S7 Fig), though 15 overlapped (*CG17839, CG32506, CG33110, Gmap, Mbs, Pdp1, Rab30, VAChT, aru, bves, fry, mip40, mxt, oys, sdk*). The relatively low overlap between these two gene sets is to be expected, since they prioritize variants of large effect vs. whole genes of moderate effect. Overall, genes annotations were enriched for neural development and function (*aru, bchs, CG13506, ChAT, Ccn, daw, dsf, Dip-δ, dpr6, fry, fz2, Mbs, Pdp1, sdk*), wing and development (*CycE, daw, dsx, egr, fry, fz2, Gart, HnRNP-K, Mbs, sno*), Rab GTPase activity (*ca, CG32506, Gmap, plx, Rab30*), and regulators of transcription (*dsf, fry, HBS1, luna, MED23, mip40, Pdp1, Rab30, SAP130, Tgi*). Different sex-based phenotypes varied in how unique certain whole genes were to a given phenotype (Fig 3C). Genes identified in the sex-average analysis were generally shared with the male and female phenotypes, while genes in the sex-difference analysis were generally unique. Interestingly, *Ccn* was present in both the male and sex-difference, and *dsf* and *sdk* were both present in the sex-average and sex-difference.

**Fig 3.**
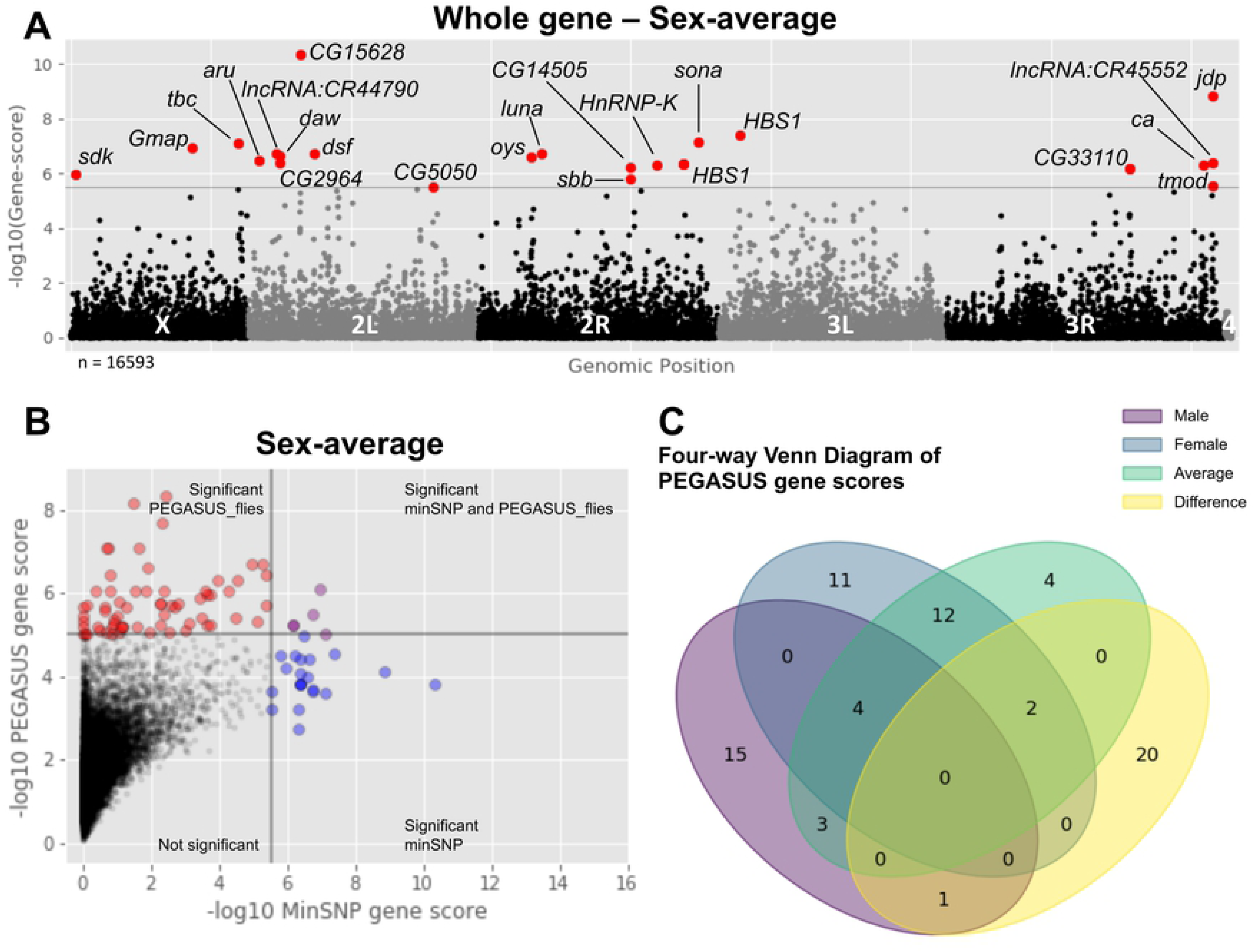
PEGASUS_flies identifies different genetic modifiers than the additive screen. (A) `PEGASUS_flies` results plotted as a Manhattan plot. For the sex-average phenotype, several genes (red points, labeled with gene symbol) exceed a strict Bonferroni significance threshold (gray dashed line, *P* ≤ 3.43E-6) identified several genes. (B) `PEGASUS_flies` prioritizes genetic modifiers of moderate effect, taking into account linkage blocks and gene length. Significant `PEGASUS_flies` (red) compared against genes significant under a minSNP approach for additive variants (blue) have very little overlap between the two sets (purple). (C) Many of the genes `PEGASUS_flies` identifies are unique to a sex-based phenotype, though the sex-average genes were generally found in other analyses.

Taking advantage of the gene-level significance scores, we leveraged publicly available gene-gene and protein-protein interaction networks to identify altered sub-networks of genes that connect to the flight performance phenotype. A local False Discovery Rate (lFDR) was calculated for each sex-based phenotype (S8 Table), for which gene-scores were either –log10 transformed if they passed or set to 0 if they did not. Transformed scores for each sex-based phenotype were analyzed together in `Hierarchical HotNet` [52], which returned a consensus network consisting of nine sub-networks of genes (S9 Table). The largest network identified 512 genes and was significantly enriched for several GO terms, including transcription factor binding, histone and chromatin modification, regulation of nervous system development, and regulation of apoptosis (S10 Table). The other eight networks were comprised of 27 genes, which together had several significant GO terms, including regulation of gene expression through alternative splicing, maintenance of the intestinal epithelium, and the Atg1/ULK1 kinase complex (S11 Table).

### Association of epistatic interactions with flight performance

Epistatic interactions account for a substantial fraction of genetic variation in complex traits [53] but they are statistically and computationally challenging to identify. To circumvent the barriers associated with performing an exhaustive, pairwise search across all possible combinations (n = 1.81E12), we turned to MArginal ePIstasis Test (`MAPIT`) to focus the search area. `MAPIT` is a linear mixed modeling approach that identifies variants more likely to have an effect on other variants. These putative hub variants represent more central and interconnected genes in a larger genetic network proposed by the Omnigenic Inheritance model [54, 55]. Accordingly, we identified 70 unique significant marginal variants exceeding a Bonferroni threshold (*P* ≤ 2.56E-8) across male, female, and sex-average phenotypes. The sex-difference analysis yielded no significant variants (S8 Fig; S12 Table). From these, only 14 had significant epistatic interactions with other variants in the genome (S13 Table), which we will discuss in order of the male, female, and sex-average results and contextualized with their epistatic interactions.

In males, there were seven significant marginal variants that mapped to five genes (*CG5645, CG18507, cv-c, sog, Ten-a*). Of the variants, only one (X_15527230_SNP) that mapped to a novel transcription start site in the BMP antagonist of *short gastrulation* (*sog*; human ortholog of *CHRD*) had significant interactions. This marginal SNP interacted with 42 other variants across 28 unique genes (S13 Table). Several of these genes are important in neuron development, signaling, and function (*CG13579*, *Dh31*, *nAChRalpha4*, *Sdc simj*, *sqz*, and *trio*), supporting accumulating evidence of a neurodevelopmental basis for variation in flight performance.

In females, there were 14 significant marginal variants that mapped to six genes (*CG6123, CG7573, CG42741, ppk23, Src64B, twi*). Of these variants, five mapped to two genes (*CG42671* and *ppk23*) with epistatic interactions. One intronic SNP (3L_11217593_SNP) mapped to *CG42671.* Little is known about this gene and there are no human orthologs, but we can gain insights into its function based on the 51 epistatic variants that mapped to 37 genes with annotations for regulation of gene expression (*arx*, *bi*, *CG6843*, *Ches-1-like*, *dve*, *HDAC1*, *Moe*, and *RpL26, Sdc*, *Tgi*), and neural development, signaling, and function (*cact, CG13579, HDAC1, ed, ngl3, nrm, numb, Sdc*). The other four variants (X_17459818_SNP, X_17459830_SNP, X_17460743_DEL, X_17460820_SNP) mapped to a 1002 bp region downstream of *pickpocket 23* (*ppk23*; human homologs in ASIC gene family). *ppk23* is a member of the degenerin (DEG)/epithial Na+ channel (ENaC) gene family that functions as subunits of non-voltage gated, amiloride-sensitive cation channels. It is involved in chemo- and mechanosensation, typically in the context of foraging, pheromone detection, and courtship behaviors [56, 57]. These marginal variants significantly interacted with 2162 variants, which mapped to 1042 genes that were also largely found in the sex-average analysis.

The sex-average phenotype had 62 significant marginal variants (11 also found in females) mapping to 21 genes (*Art2, CG10936, CG15630, CG15651, CG18507, CG3921, CG42671, CG42741, CG5645, CG6123, CG9313, CR44176, cv-c, Fad2, natalisin, ppk23, Rbfox1, Rgk1, Src64B, twi*). Of the 62 marginal variants, 18 had significant epistatic interactions: nine were intergenic, seven mapped to *ppk23*, and the remaining four mapped to single genes: *CG42671*, *CG10936, CG9313*, and *CG15651* (S13 Table). Previously identified in the female analysis, *ppk23* had the greatest number of interactions, placing it close to the center of a highly interconnected genetic landscape (Fig 4A). The seven marginal variants interacted with 4895 variants across 2010 unique genes, 11 of which mapped to genes that also contained significant marginal variants (*A2bp1, cv-c, Fad2, CG9313, CG10936, CG42741, Rgk1, sog, Src64B, twi, Ten-a*). The 2010 unique genes had significant GO term enrichment for neuronal growth, organization and differentiation (S14 Table). One of *ppk23*’s interactors was *CG42671,* itself a gene with a significant marginal variant in the sex-average epistasis screen and previously mentioned in the female epistasis screen. For the sex-average epistasis screen, *CG42671* interacted with 1013 variants across 616 genes. These genes were significantly enriched in a gene set enrichment analysis for genes involved in neurodevelopment, particularly neuron growth and movement (S15 Table). While this gene is understudied and lacks substantive annotations, but based on its interactors’ significant GO categories, it is very likely CG42671 is involved in growth and neuronal target finding. *CG10936* has few annotations, though it was identified in a screen for nociception [58]. It paired with 29 genes annotated for neurogenesis and function (*CG42788*, *Dh31*, *fru, hiw, lilli*, *nAChRalpha4*), as well as regulation of gene expression through chromatin modification (*Etl1* and *lilli*) and alternative splicing (*Srp54* and *U2af38*). One SNP (2R_16871314_SNP) was mapped to both the 3’ UTR of *CG9313* and 29 bp downstream of *CG15651*. *CG9313* (orthologous to human DNAI1) is an ATP-dependent microtubule motor and is involved in the sensory perception of sound in *Drosophila* and proprioception, as well as sperm development [59]. *CG15651* is predicted to localize to the rough endoplasmic reticulum and Golgi body during embryogenesis, early larval, and late pupation stages where it is expressed in the central nervous system. Its human ortholog, FKRP (fukutin related protein), is implicated in intellectual disability and it is a candidate gene therapy target for muscular dystrophy [60–62]. These genes’ shared function in nervous system development is reflected in the variants that map to 87 genes with annotations for neuron development, patterning, and function (*5-HT2B, cwo, dally, dx, Dysb*, *enok*, *erm, mbl, Ngl1, nmo, Sdc, Sema1a, sNPF*, *tup*,). Several genes were also annotated for endoplasmic reticulum function (*bark*, *CG5885*, *CG15651*, *Fatp3*, *PAPLA1*, *Trc8*, *Uggt*); chromatin remodeling (*CG43902, enok, erm, lncRNA:roX1*, *tim*); transcription and alternative splicing (*cwo, bru3, CG6841, CG9650, CG15710, enok, luna, mbl, tim, tup*); and gene product regulation (*bru3, cwo, CG5885, CG9650, CG15710, luna, tRNA:Arg-TCT-2-1, tup*). Finally, there was a 669 bp region with six intergenic variants (chr3L:6890373 - 6891042). This region lacked regulatory annotations, yet collectively interacted with 513 variants mapping to 309 genes, many of which were shared with *ppk23*, *CG42671*, and *CG10936*. Similarly, these genes had significant GO term enrichment for neurodevelopment and neuron function (S16 Table).

**Fig 4.**
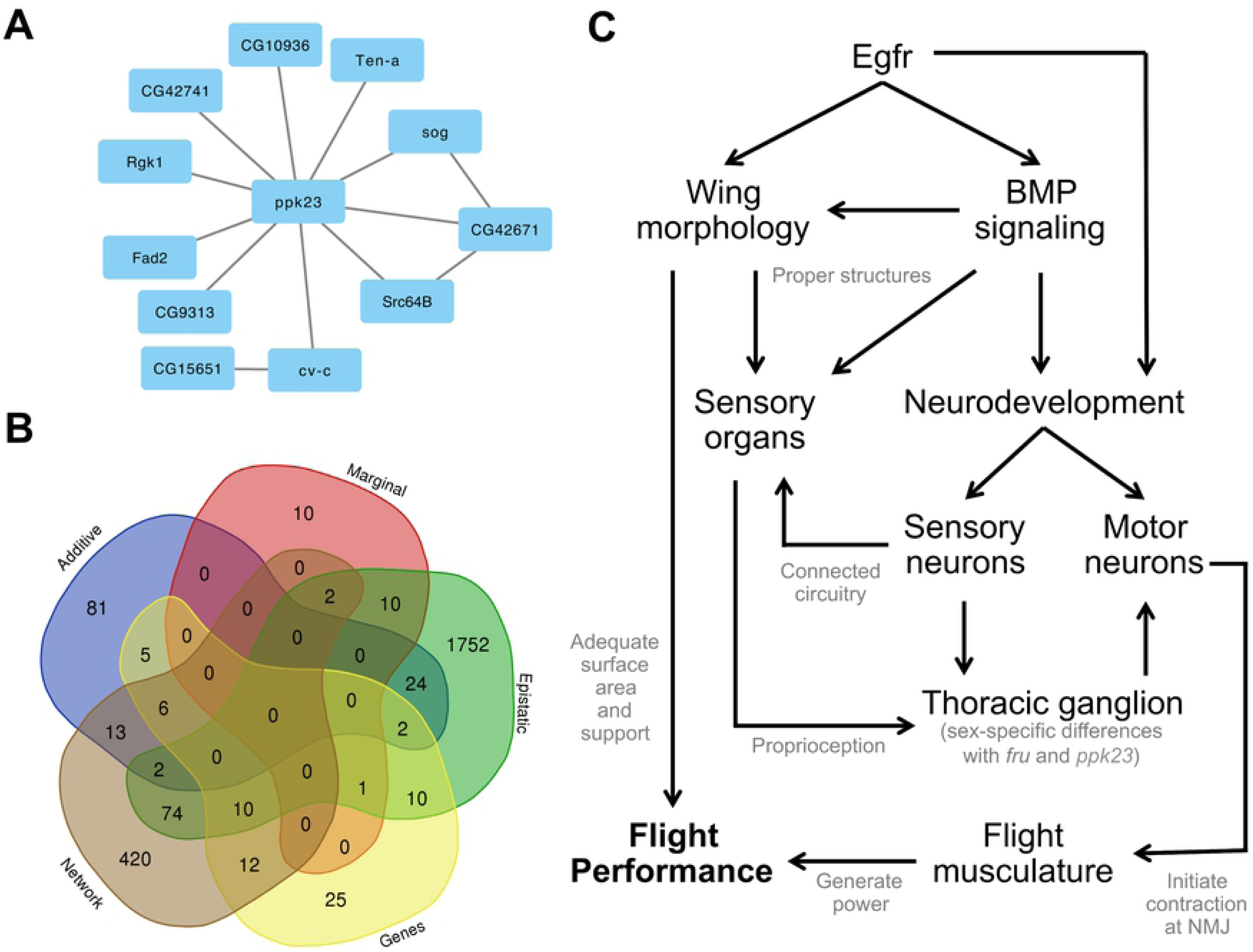
Flight performance is a larger complex trait comprised of several smaller traits. (A) The genetic architecture of epistatically interacting genes generally coordinated through *ppk23*. A few other genes mapped to from marginal variants had epistatic interactions with marginal variants in *ppk23*. (B) Genes or genes mapped to from variants across different analyses were not identified in more than three analyses. Roughly half or more genes were unique to each analysis. (C) Fight performance has a complex genetic architecture, with the key developmental gene *Egfr* and BMP signaling pathway contributing to wing and neurodevelopment. These processes are both important for structuring the sensory organs that enable the fly to use mechanosensory channels for proprioception. Signals from the sensory organs on the wing, head, and body travel to the brain and thoracic ganglion, which sends signals through the motor neurons to the direct and indirect flight musculature that is also differentially assembled and innervated to generate power and control the wing angle during flight.

There were epistatic interactions between several of the genes identified from marginal variants (Fig 4A). Since marginal variants represent those more likely to interact with other variants, their interaction with one another suggests a highly interconnected genetic architecture underlying flight performance. Additionally, the breadth of epistatic interactions from a small, focused subset of marginal variants supports an important role for epistasis in the genetic architecture of flight performance. There are likely many more variants that interact with one another. But based on strong enrichment for neurodevelopmental genes from the very limited subset of marginal variants we tested, we hypothesize that flight performance in wildtype *Drosophila* is modulated by neural function and circuitry.

### No evidence for adult transcriptome variation affecting flight performance

Since many variants mapped to cis-regulatory elements and trans-regulatory genes, we sought to test whether regulatory variation was affecting developmental or adult homeostasis. Accordingly, performed a Weighted Gene Co-expression Network Analysis (WGCNA)[63] using 177 publically available DGRP transcriptomic profiles for young adults of both sexes [64]. We clustered genes by similarity in expression profile, then correlated those clusters’ eigenvalues with the mean and standard deviation of flight performance, as well as the proportion of flies that fell through the column over the total assayed. No clusters across sex or phenotype had a significant correlation. This result squares well with our previous observation that many of the significant variants map to genes involved in pre-adult development, rather than genes that maintain adult homeostasis (S9 Fig).

### Flight performance is modulated by an interconnected genetic architecture

The genetic architecture of flight performance is comprised of many different types of genetic modifiers. Many of the variants map to genes that are found across analytic platforms (Fig 4B). Most variants were unique to a single analysis, suggesting that association studies should consider using multiple different analyses to enhance the power to detect variants and genes in their study. However, many genes and genes identified from variants were identified in two (148) or three (23) analyses. Those involved in three analyses include: *aru, CG2964, CG13506, CG15651, CG17839, CG42671, CycE, daw, Diap1, Egfr, fz2, Gart, Gmap, Mbs, MED23, mip40, mxt, Pdp1, Rab30, rhea, sog, sona, Tgi*) analyses. This suggests that individual genes can contain variants with different types of effects or have differential contributions to the overall genetic architecture. A complete lookup table of all genes and genes identified from variants is available (S17 Table).

## Discussion

We tested flight performance of 197 DGRP lines, identifying several additive and marginal variants, epistatic interactions, whole genes, and a consensus network of altered sub-networks that associated with variation our phenotype. We identified many cis-regulatory variants mapped to genes with annotations for wing morphology, indirect flight muscle performance, and neurodevelopment of sensory and neuromuscular junctions.

### Variation in gene regulation drives variation in flight performance

Variation in gene expression is a major contributor to phenotypic variation [65, 66]. Association studies with the DGRP lines often map variants to intergenic and non-coding regions of genes [22, 48, 67]. These regulatory elements can be cis-regulators, like transcription factor binding sites (TFBS), enhancers, or silencers; or they can be trans-regulatory, like transcription factors, splicosomes, or chromatin modifiers. In the present study, the vast majority of variants in the additive, marginal, and epistatic analyses mapped to introns or within 1kb of a gene, suggesting a cis-regulatory role.

When cis-regulatory elements lie in important developmental genes, their effects can be magnified as the organism continues through development. The most significant additive variant we identified mapped to an *epidermal growth factor receptor* (*Egfr*; human homolog *EGFR*) intron. Encoding a key transmembrane tyrosine kinase receptor, Egfr is a pleiotropic gene affecting developmental and homeostatic processes throughout the life and anatomy of the fly. It is well known for its role in embryonic patterning and implications in human cancers [68, 69]. The variant also mapped to several overlapping TFBS for transcription factors known to affect embryonic development in a highly dose-dependent manner (*bcd, da, dl, gt, hb, kni, Med, prd, sna, tll, twi, disco, Trl*). Variation in patterning cells fated to become tissues and organs can be magnified through the adult stage, especially when that receptor is also known to affect other developmental processes [70]. Other intronic variants were identified in *Egfr* through the epistatic interactions with *ppk23*, illustrating how different types of genetic modifiers can exist within the same gene.

The role of cis- and trans-regulatory elements goes even further when there is variation in cis-regulatory elements of trans-regulatory genes. One of the Bonferroni additive variants mapped to an intronic region of *Forkhead domain 96Ca* (*fd96Ca*; human homologs *FOXB1* and *FOXB2*), a TFBS for *dorsal (dl)*, and a silencer for *histone deacetylase 1* (*HDAC1*). *fd96Ca* is a fork head box transcription factor expressed in neuroblasts along the longitudinal axis of the embryo and in some sensory neurons in the embryonic head [71]. Trans-regulators, like fd96Ca, are proposed to have a large impact on phenotypic variation under the Omnigenic Inheritance model [54, 55]. Similar to *Egfr*, regulatory variation in a gene that helps determine cell fates can have larger effects if not enough cells are allocated for differentiation later in life. This can begin a cascade that amplifies downstream [72] and may hint at why trans-regulators were significant Gene Ontology (GO) terms in the consensus network.

There are likely non-coding regions of the genome that correspond with more cryptic regulatory regions. Six intergenic, marginal variants in a 669bp stretch (chr3L:6890373 - 6891042) had a number of significant epistatic interactions with developmental and neurodevelopmental genes. These variants lacked regulatory annotations in the DGRP2 annotation file, however these annotations were collected during embryogenesis [47] so it is possible these sites are activated by trans-regulators during different times in development. Nonetheless, based on its epistatic interactions, it is likely an important cis-regulatory region that affects general development from an early stage in the fly life cycle.

Our results suggest genetic variation in regulatory (non-coding) regions has a greater affect on variation of flight performance than variation in protein coding regions. While non-synonymous variants can have large effects on flight performance [73–75], they were uncommon in our screen compared with variation in non-coding regions. This may be a result of strong purifying selection acting against them in a natural setting. Many of the candidate modifiers of flight are more commonly expressed during development [39, 45, 46]. This observation is supported by our lack of evidence for adult transcriptomic variation correlating with flight performance. Additionally, the flight phenotype was highly heritable, suggesting our phenotype was not an artifact of environmental or experimental variation. Finally, the constructs we used to validate candidate genes created genetic variation in intronic regions, rather than post-transcriptionally modifying gene expression with an RNA^i^ construct. Our successful validation of several candidate genes suggests variation in the non-coding regions of the candidate genes is sufficient for observing phenotypic differences. Further, insertion of the constructs into intronic regions both positively and negatively affected performance, even when done at independent sites in the same gene, suggesting a more nuanced impact of genetic variation in cis-regulatory regions. We conclude that modifiers of cis- and trans-regulation in pre-adult stages are more likely to modify flight performance in wild populations than variation in coding sequence.

### Variation in wing and indirect flight muscle development contributes to variance in flight performance

Flight performance is a complex trait comprised of coordination across several smaller developmental and functional, complex traits[13, 76, 77]. The central role of *Egfr* in development means it can have wide range of functional effects on adult morphology. Natural variants in *Egfr* are known to cause developmental differences in wing morphology that can significantly alter flight performance [70, 77], in part through interactions with the Bone Morphogenetic Protein (BMP) signaling pathway [13, 70, 78]. BMP signaling is also an established modifier of wing development, as it forms dose-dependent gradients that pattern the wing size and shape [79, 80], as well as sensory and neuromuscular circuits [6, 81]. We identified several modifiers of BMP signaling (cmpy, Cul2, *cv-2, cv-c, dpp, dally, daw, egr, gbb, hiw, kek5, Lis-1, Lpt, lqf, ltl, Mad, nmo, scw, srw, Snoo, tkv, trio*) across all analyses and functionally validated *Snoo*— discussed below. Among the modifiers of BMP signaling, *short gastrulation* (*sog*; human homolog *Chordin*) stood out as a known source of natural variants that modifies flight performance in natural populations [13]. *sog* affects wing morphology through its role as a *dpp* antagonist in patterning the dorsoventral axis of the wings [80, 82, 83]. *sog* is also noteworthy for its interconnectedness to other genes containing both a significant marginal variant and variants that had epistatic interactions with other significant marginal variants: *ppk23* and *CG42671* (formerly *CG18490* and *CG34240*)—discussed below. Marginal variants represent a class of variants that are statistically more likely to interact with other variants [84], via epistasis. Their identification hints at a more interconnected role in the genetic architecture. In this case, identification of *sog* suggests a more interconnected role for this antagonist of BMP signaling in modifying flight performance.

In addition to wing morphology, we identified several modifiers known to affect flight muscle function. The indirect flight muscles (IFM) power flight through the alternating dorsoventral and dorsolongitudinal muscle contraction to deform the cuticle and move the wings [85, 86], while the direct flight muscles control flight through precise adjust of the wing angle [87]. We identified two genes with known roles in flight [88, 89] from the additive screen that we successfully validated: *Lasp* and *bru1*. *Lasp* (human ortholog *LASP1*), is the only nebulin family gene in *Drosophila*, and shown to modify sarcomere and thin filament length, and myofibril diameter [88]. We also identified *bruno 1* (*bru1* or *aret*; human homolog *CLEF1* and *CLEF2*), a transcription factor that controls alternative splicing of myofibrils in the IFM [9, 89], among other developmental processes. *bru1* had two intronic variants, one of which mapped to a TFBS for *twi*—one of the genes identified from a significant marginal variant.

Using our newly developed platform `PEGASUS_flies` to find significant whole genes, we also identified *tropomodulin* (*tmod;* human homolog *TMOD1*) and *Glycerol-3-phosphate dehydrogenase 1* (*Gpdh1*; human homolog *GPD1*). These two genes were previously validated for their roles in flight performance [8, 90] and are responsible for muscle function and metabolism within muscles, respectively. The identification and previous validation of *tmod* and *Gpdh1* is noteworthy because neither had a significant variant exceed the additive screen’s significance threshold (*P* ≤ 1E-5). This finding demonstrates a successful proof-of-principle for `PEGASUS_flies`’ ability to identify genetic modifiers that would otherwise be overlooked in a traditional minSNP approach in an additive screen. Additionally, we successfully validated *fry*, identified in both the additive and whole gene screens. Taken together, the prior and current validation of these genes establishes `PEGASUS_flies` as a verified platform for identifying modifiers of complex traits.

### Neurodevelopmental genes play an important role in modifying flight performance

Many neurodevelopmental genes with diverse functions were identified across analyses. Because neurodevelopmental genes can play several roles, many of which are unannotated in GO databases, GO term enrichment analyses can be underpowered. This may explain why we failed to identify any GO terms for additive variants in the GOwinda analysis [22]. However, their identification through other GO analyses on the epistatic and network-based analyses is encouraging.

Several neurodevelopmental genes overlapped between the additive minSNP and PEGASUS_flies whole gene approach. These genes (*aru, ChAT, Ccn, DIP-δ, dsf, dsx, fry, Mbs, sdk, VAChT*), lend additional support to the likelihood these genes were not false positives. For example, *fry* and *Sidekick* (*sdk*) both coordinate dendritic target finding functions with DSCAM family genes [91, 92]. This is in agreement with several significant GO terms for axon guidance and neuronal targeting in the consensus network’s largest sub-network (S11 Table) and for the genes identified from epistatic interactions with *ppk23*, *CG42671*, and an intergenic region (chr3L:6890373 - 6891042)(S14-16 Tables). Accordingly neurodevelopmental genes are present throughout our study, and represent a highly interconnected group of genes that likely plays an important role in flight performance.

Underscoring this interconnectedness is the identification of several neurodevelopmental genes that mapped to epistatic interactions with a common, significant marginal variant in *sog*. This variant was significant in males and mapped to a new transcription start site. In addition to affecting wing morphology, *sog* also plays a role in neurodevelopment (*CG13579, dib, Hk, lncRNA:rox1, nAChRα4, Sdc, simj, sqz, Toll-4, trio*) [36, 37, 81, 83]. Several of these genes were involved in neuromuscular growth and function (*CG13579, Hyperkinetic (Hk), nicotinic acetylcholine receptor α 4 (nAChRα4), Syndecan (Sdc), squeeze (sqz), trio*) [81, 93–97], suggesting an important connection between neurodevelopmental phenotypes and their role in activating direct and indirect flight muscles. However, some of the genes interacting with *sog* can affect sensory neurons as well. For example, *trio* is also present in sensory neurons and is capable of modifying chemosensation [21]. Other *sog* variants that had epistatic interactions with marginal variants in *CG42671* (formerly *CG18490* and *CG34240*) and *ppk23*—discussed below, two genes with known or putative roles in developing the peripheral nervous system (PNS).

In addition to neuromuscular genes, we validated genes involved in patterning the PNS. One of the Bonferroni variants from the additive screen mapped to *Down Syndrome Cell Adhesion Molecule 4* (*Dscam4*; human ortholog *DSCAM*). DSCAMs are a conserved family of extracellular, immunoglobin proteins that promote cell-cell adhesion. They are found in complex (type IV) dendrite arborization neurons that promote dendritic target recognition and dendrite self-avoidance in the developing PNS [34] and in the brain and central nervous system (CNS) [98, 99]. Type IV dendritic arborization neurons transduce signals from sensory neurons (e.g. photoreceptors, chemosensors, and mechanosensors), to the CNS [99–102]. Dscams are expressed differentially and combinatorally in different neurons, which allows them to create highly interconnected neural circuits [99]. They also work with other cell-cell adhesion proteins, like cadherins, in patterning the nervous system. *Cadherin-N* (*CadN* or *N-cad*) interacts with *Dscam2* and *Dscam4* in patterning olfactory receptor neurons (ORN), like *Or46a* (significant additive hit) and *Or59c* (significant epistatic hit with *ppk23*) [102–105]. Given their importance in patterning sensory neuron circuits and strong significance in the additive screen, we independently validated *Dscam4* and *CadN* using two separate insertional mutants for each. Both pairs of insertional mutants in both genes were significant, though the direction of effect was reversed, reiterating how cis-regulatory regions can differentially affect genes’ expression levels. Our double validation for each supports a greater level of confidence in *Dscam4* and *CadN* as modifiers of the peripheral nervous system important for flight performance.

We validated two other dendrite patterning genes that also help to form sensory organs on the wing and body that contribute to proprioception: *furry* (*fry*; human homolog FRYL) and *Sno oncogene* (*Snoo or dSno*; human homolog SKI). These two conserved proteins are expressed along the same types of sensory neurons as Dscams and cadherins that promote dendrite field patterning, dendrite self-avoidance, and development of sensory organs [106]. *fry* assists Dscams and cadherins in dendritic tiling of chemosensors (olfaction or gustation) and mechanosensors (proprioception) [105–107] that directly connect to sensory microchaete (hairs or bristles) organized along the wing and body in specific patterns [108]. Meanwhile, *Snoo* interacts with the wingless pathway [6, 109], and is an important antagonist of *Medea* (*Med* or *dSmad4;* human homolog *Smad4*)—an important regulatory of the BMP-to-activin–β pathway [110]. *Snoo* is known to modify wing shape [110], dendritic tiling, and the development of sensory organs (microchate and campaniform sensilla) on the wing [6, 111]. These sensory organs play different roles; wing chaete can function as chemosensors (olfaction and gustation) and mechanosensors [100, 112], while campaniform sensilla measure strain on the deformed wing blade [113–116]. Together, these sensory organs aid in proprioception of flight [11] and delineate a direct connection between the role of proper development of the wings’ sensory organs and the proper development of the neural circuitry connecting them to the CNS in modifying flight performance.

We functionally validated two candidate genes with only tangential evidence of their function that we are naming *flapper* (*flap*, formerly *CG11073*) and *flippy* (*flip*, formerly *CG9766*). *flapper* is expressed in the peripodial epithelium cells of the eye, leg, and wing imaginal discs [117]. It is expressed at very high levels during 16-18 hours of embryogenesis, pupariation [45] and in the head, eyes, and carcass in the adult stage [118]. It was previously identified as a candidate gene in a screen for modifiers of circadian rhythm [119] and was significantly upregulated in flies bred for aggressive behavior [120], but both studies failed to functionally validate the gene. *flapper* was also implicated in the downregulation of amyloid-β peptides [121] and in late life fecundity [122] suggesting it may play a basic role in development that affects several phenotypes. Accordingly, we hypothesize it plays some role in patterning neural circuitry of sensory neurons on the cuticle and eyes, and facilitates neural circuit assembly in the brain. The other gene, *flippy* (human homolog *FANK1*), is pleiotropic with important roles in neuroanatomical development [43, 123] and sperm development [46]. It is important in the development of trichogen cells, which are precursors to the chaete flies use for mechanosensation. In humans, *FANK1* plays roles in spermatogenesis and apoptosis, and is a putative evolutionary target of balancing selection [124, 125]. Given *flippy*’s pleiotropic role in neurodevelopment and gametogenesis, it may also be under stabilizing selection brought about by contrasting selective pressures for neural function and fitness.

Finally, qualitative observations of differentially performing DGRP lines support a role for proprioception as a modifier of flight performance. High-speed videos of strong, intermediate, and weak lines show strong lines react quicker to an abrupt free fall and are better at controlling their descent than the intermediate fliers, and much more than weak fliers. This direct evidence corroborates with the validation screen and inferential association analyses to support a role for natural variants in genes that affect 1) sensory neural circuit connectivity, 2) development and function of neuromuscular junctions, and the integration of these two onto wings of varying morphologies for modifying flight performance in a natural population.

### Important implications for acid sensing ion channels in flight performance and neural function flight

Pickpocket genes encode a conserved group of degenerin/epithelial sodium channels (DEG/ENaC) that function as non-voltage gated, amiloride-sensitive cation channels [56]. They are found in the brain, thoracic ganglion [57, 126], neuromuscular junctions[126, 127], and trachea [128], though pickpocket family genes are most commonly found along type IV dendrite arborization sensory neurons that connect chemo- or mechanosensory organs to the CNS [105-107, 126, 129-132] on the head, legs, and wings [57, 127, 133–136]. Chemosensing microchaete can contain olfactory receptor neurons (ORN), gustatory receptor neurons (GRN), and ionotropic receptors (IR), which are useful for foraging and pheromone detection [11, 57, 127, 133–139]. In this study, we identified six pickpocket genes (*ppk1, ppk8, ppk9, ppk10, ppk12, ppk23*), 10 gustatory receptors (*Gr10a, Gr10b, Gr28b, Gr36b, Gr36c, Gr39a, Gr59a, Gr59d, Gr61a, Gr64a),* 12 olfactory receptors and binding proteins (*Or24a, Or45a, Or46a, Or49a, Or59b, Or59c, Or67d, Or71a, Or85d, Obp8a, Obp28a, Obp47a),* and 13 ionotropic receptors (*Ir41a, Ir47a, Ir47b, Ir51a, Ir56b, Ir56c, Ir56d, Ir60d, Ir60f, Ir62a, Ir64a, Ir67b, Ir75d)* from the additive, marginal, epistatic, and network approaches. *Or85d* was identified from the 2^nd^ most significant additive variant and only non-synonymous SNP that passed a Bonferroni threshold in the additive search. And yet, despite a combined 41 pickpocket, gustatory receptor, olfactory receptor, and ionotropic receptor genes, only six (*ppk10*, *ppk12*, *Gr59d*, *Or24a*, *Ir41a*, and *Ir60d*) overlapped with an olfactory screen testing for genetic associations across 14 odors [21]. Accordingly, we hypothesize a more nuanced role for these chemosensors in aiding proprioception during flight.

The magnitude of significant marginal variants and epistatic interactions that mapped to *ppk23* suggests this ion transporter has a much more interconnected role in the genetic architecture of flight performance than previously thought. *ppk23* is a modifier of flies’ ability to track odors during free flight, but not a modifier of odorless flight [140]. Our results support a role for *ppk23* in modifying flight, along with all but eight (*Or46a, Or49a, Or85d, Gr36b, Gr36c, Ir60d, Ir60f, ppk10*) of the 41 previously listed pickpocket and chemoreceptor genes that *ppk23* interacted with. Like *sog*, *ppk23* is likely a central modifier of performance based on the number of epistatic interactions with variants mapping to genes identified in the marginal variant screen (*A2bp1/ Rbfox1, cv-c, Fad2, CG9313, CG10936, CG42741, Rgk1, sog, Src64B, twi, Ten-a*). Some of these play roles in sensory signal processing (*A2bp1/ Rbfox1, CG9313*, *CG10936*, *Fad2*, *Rgk1*), neuron growth (*sog* and *Src64b*), neuromuscular junction development (*cv-c*, *Src64b*, *Ten-a*), and transcription factors (*A2bp1/ Rbfox1, CG42741*, *twi*) [141, 142], several of which had significant epistatic interactions of their own. Of these, *CG10936* is proposed to be involved in sensory perception [142], but has limited annotations otherwise. Our work supports this hypothesized function. *ppk23,* in addition to these interactions, is known to modulate physiology and lifespan [143], broadening its canonical roles in chemo- and mechanosensation. Taken together, *ppk23* likely has strong connections to many systems beyond detection of stimulation that have deeper connections to organismal biology.

The interconnectedness of *ppk23* also provides clues about the sexual dimorphism observed in flight. While males generally outperform females, likely due to differences in weight, sex failed to explain ∼25% of the variation between the two groups. Like most pickpocket family genes, *ppk23* is well established as an important factor in chemosensation, pheromone detection, and courtship [57, 126, 130]—highly sex-specific phenotypes. One of *ppk23*’s epistatic interactions mapped to *fruitless* (*fru*; human homolog *ZBTB24*), a transcription factor responsible for sex-specific neural phenotypes involved in courtship and pheromone detection [144] that co-localizes with *ppk23* differentially between sexes, on the leg and wing microchaete [4, 57, 126, 127, 143]. In addition to the PNS, *ppk23* and *fru* have sex-specific co-localization patterns in the thoracic ganglion. This cluster of neurons central to the “escape” response, allowing for ultra-fast processing of and response to flight-associated cues [11, 145]. Males show more connections between *ppk23* and *fru* in the thoracic ganglion, and co-localization in neurons crossing the midline between the two sides of the anterior-most, pro-thoracic ganglion [57, 126]. *fru* is also expressed in vMS2 motor neurons connecting the thoracic ganglion to the flight musculature, likely involved in courtship song generation and aggression behaviors [146, 147]. The connection between sensory neurons, *ppk23*, *fru*, and motor neurons involved in wing motion draw a clear connection between a potential mechanism delineating the sex-difference phenotype we observed. Given the prior connections between *ppk23*, *sog*, and the epistatic interactions between them that annotate to sensory neurons and motor neuron neuromuscular junctions, there are likely other important connections underlying the ability of flies to process proprioceptive signals that are relayed directly to the flight musculature during our assay that have yet to be uncovered. Some of these connections may lie in the genes identified using `PEGASUS_flies`’ for the sex-difference analysis, like *doublesex* (*dsx*), an interactor of *fru* and *ppk1* in patterning sex-specific neural networks for courtship; dissatisfaction (*dsf*), a modifier of courtship behavior [146, 148–150]; and several other genes: *blue cheese (bchs), Ccn, CG13506, defective proboscis extension response 6 (dpr6), pollux (plx), sidekick (sdk), eiger (egr*) [4, 34, 151, 152]. Further study of these genes may yield promising insights into the sex-differences we observed in flight performance, as well as sex-specific behavioral traits.

### A proposed model for understanding the genetic architecture of flight performance

Flight performance is likely an epiphenomenon of several interconnected complex traits. While we are unable to identify every modifier, we likely identified the main components of the genetic architecture. Accordingly, we propose the following model to synthesize our findings (Fig 4C).

Epidermal growth factor receptor is a key gene in a canonical developmental pathway. It can affect wing morphology, sensory organ development, and neurodevelopment, on its own and through the BMP signaling pathway. Proper development of these structures and circuits enables well-connected sensory neurons to receive external stimuli regarding proprioception. These signals are transduced through the thoracic ganglion, with sex-specific differences potentially modulated through *ppk23*, *fru*, and *dsx*. The thoracic ganglion processes these signals and activates motor neurons, which innervate the direct (control) and indirect (power) flight musculature at neuron muscular junctions. Activating these muscles allows the properly developed wings to flap and generate lift.

### Implications for BMP signaling and pickpocket genes in neuroinjury and neurodegeneration

The complexity of congenital, neurodegenerative diseases lies in the mix of genetic elements with very modest effect size. Association screens with *Drosophila* present a compelling model for identifying these sources of variation, especially in neuron-centric traits [25, 48, 153, 154]. Our results present a strong link between flight performance and BMP signaling—a proposed candidate pathway for therapeutic interventions in several neurodegenerative diseases [30, 32, 155]. Mutations in *thickveins* (*tkv*) human homologs *BMPR1A* and *BMPR1B* are linked to familial Alzheimer’s Disease [156], while mutants of *Superoxide dismutase 1* (*dSOD1*; human homolog *SOD1*) associated with Amyotrophic Lateral Sclerosis (ALS) can be rescued by activators of BMP signaling expressed in proprioceptive and motor neurons [157]. Our validation of the BMP antagonist *Snoo* confirms BMP signaling plays a role in flight performance. Given the number of epistatic interactions between *ppk23* and BMP signaling genes, it is very likely our data uncovers important modifiers of the BMP pathway that affect neurodysfunction in humans.

In addition to BMP signaling, we propose an expanded role for ppk23, and pickpocket family genes more generally, in neurobiology and neurodysfunction therapeutics. Acid Sensing Ion Channel (ASIC) family genes, the human homolog of the pickpocket family, can function as neuronal damage sensors. They detect drops in pH around neurons, often caused injury, damage, and dysfunction, which can elicit an inflammation response [31, 158]. These channels are found all over the brain and spinal column, supporting a functional and protective role following traumatic brain injury (concussion) and cerebral ischemia (stroke) [29, 31]. They are also identified as a potential target for genetic and/or pharmacological interventions of neurodegeneration and neuroinflammation [158]. Accordingly, our results break ground in identifying candidate genetic interactions that might be useful for such interventions.

## Materials and Methods

### Drosophila Stocks and Husbandry

All stocks were obtained from Bloomington *Drosophila* Stock Center (https://bdsc.indiana.edu/), including 197 *Drosophila* Genetic Reference Panel (DGRP) lines [20], 23 *Drosophila* Gene Disruption Project lines using the Mi{ET1} construct [159, 160], and two genetic background lines (w^1118^ and y^1^w^67c23^; S1 Table).

Flies were reared at 25° under a 12-h light-dark cycle. Stocks were density controlled and grown on a standard cornmeal media [161]. Two to three days post-eclosion, flies were sorted by sex under light CO_2_ anesthesia and given five days to recover before phenotyping.

### Flight performance assay

Flight performance was measured following the protocol refined by Babcock and Ganetzky [33]. Briefly, each sex-genotype combination consisted of approximately 100 flies, divided into groups of 20 flies across five glass vials. These vials were gently tapped to draw flies down, and unplugged before a rapid inversion down a 25 cm chute. Vials stopped at the bottom, ejecting the flies into a 100 cm long x 13.5 cm diameter cylinder lined with a removable acrylic sheet coated in TangleTrap adhesive. Free falling flies instinctively right themselves before finding a place to land, which ended up immobilizing them at their respective landing height. Flies that passed through the column were caught in a pan of mineral oil and were excluded from the analysis.

After all vials in a run were released, the acrylic sheet was removed and pinned to a white poster board. A digital image was recorded on a fixed Raspberry PiCamera (V2) and the x,y coordinates of all flies were located with the ImageJ/FIJI Find Maxima function with a noise tolerance of 30 [162]. For each sex-genotype combination, the mean landing height was calculated for only the flies that landed on the acrylic sheet.

### High-speed video capture of flight column

High-speed videos of flies leaving the flight column were recorded at 1540 frames per second using a Phantom Miro m340 camera recording at a resolution of 1920 x 1080 with an exposure of 150 μs (Data available in File S1). The camera was equipped with a Nikon Micro NIKKOR (105 mm, 1:2.8D) lens and Veritas Constellation 120 light source.

### Estimating heritability

Individual fly landing heights were adjusted for covariate status by adding the difference between the DGRP webserver’s adjusted and raw line means for each sex, and added them back to the individual landing height of the respective sex and genotype. Using these adjusted landing heights by sex, we performed a random effects analysis of variance using the R (v.3.5.2) package lme4 (v.1.1.23): *Y ∼ μ + L + ε*. Here, *Y* is the adjusted flight score, *μ* is the combined mean, *L* is the line mean, and *ε* is the residual. From this, sex-specific broad sense heritability (*H^2^*) estimates were calculated from the among line (*σ_L_^2^*) and error (*σ_E_^2^*) variance components: *H* = *σ_L_^2^* / (*σ_L_^2^* + *σ_E_^2^*).

### Genome wide association mapping

Flight performance scores for males and females were submitted to the DGRP2 GWAS pipeline (http://dgrp2.gnets.ncsu.edu/) [19, 20] and results for each sex, and the average (sex-average) and difference (sex-difference) between them were all considered (S3 Table). In total, 1,901,174 variants with a minor allele frequency (MAF) ≥ 0.05 were analyzed (Data available in File S2). All reported additive variant *P*-values result from a linear mixed model analysis, including *Wolbachia* infection and presence of five major inversions as covariates. Variants were filtered for significance using the conventional *P* ≤ 1E-5 threshold [48]. Effect size estimates were calculated as one-half the difference between the mean landing heights for lines homozygous for the major vs. minor allele. The contribution of individual variants to the overall effects was estimated as the absolute value of an individual variant’s effect size divided by the sum of the absolute values for all conventionally significant (*P* < 1e-5) variants’ effect sizes.

### Candidate gene disruption screen

Candidate genes were validated using insertional mutant stocks generated from Gene Disruption Project [51]. These stocks contain a *Minos* enhancer trap construct Mi{ET1}[50] and were built on either w^1118^ or y^1^ w^67c23^ backgrounds (BDSC_6326 and BDSC_6599, respectively).

Control and experiment line genetic backgrounds were isogenized with five successive rounds of backcrossing the insertional mutant line to its respective control. Validation of flight phenotypes was done using offspring of single-pair (1M x 1F) crosses between the control and insert lines. Heterozygous flies from these crosses were mated in pairs and the homozygous offspring lacking the insertion were collected as the control. Candidate heterozygous/homozygous positive lines were mated as pairs once more and lines producing only homozygous positive offspring were used as experimental lines (S1 Fig). Experimental lines were checked for a GFP reporter three generations later to confirm their genotype. The finalized recombinant backcrossed control and experimental lines for each sex-genotype combination were assayed for flight performance, and tested for significance, via Mann-Whitney U-tests.

### Calculating gene-score significance

Gene-scores were calculated using Precise, Efficient Gene Association Score Using SNPs (`PEGASUS`) [28]. Originally implemented with human datasets, we modified the program to work with *Drosophila* datasets, which we call `PEGASUS_flies`. It also contains default values adjusted for *Drosophila*, a linkage disequilibrium file, and gene annotations drawn from the FB5.57 annotation file, available on the DGRP webserver. `PEGASUS_flies` is available at: https://github.com/ramachandran-lab/PEGASUS_flies, and as File S4.

### Identifying altered sub-networks of gene-gene and protein-protein interaction networks

Returned gene-scores were filtered for genes of high confidence using the `Twilight` package (v.1.60.0) in R (Scheid and Spang 2005). Here, we estimated the local False Discovery Rate (lFDR) of all previously output gene scores using the *twilight* function. Taking the inflection point of the (1 – lFDR) curve, our high-confidence gene scores ranged from 0.65 – 0.73 for the four, sex-based phenotypes (S8 Table). High confidence genes were –log10 transformed, while the remaining were set to 0.

`Hierarchical HotNet` was used to identify altered sub-networks of interacting genes or proteins [52] based on network topology generated from several gene-gene or protein-protein interaction networks. The four adjusted, sex-based gene-score vectors were mapped in the program to fifteen interaction networks obtained from High-quality INTeractomes (HINT)[163], the *Drosophila* Interactions Database (Droidb)[164, 165], and the *Drosophila* RNA^i^ Screening Center (DRSC) Integrative Ortholog Prediction Tool (DIOPT)[166]. Consensus networks were calculated from 100 permutations of all four gene-score vectors on each of the fifteen interaction networks and filtered to include at least three members. The largest sub-network and the remaining eight sub-networks were passed to Gene Ontology enRIchment anaLysis and visuaLizAtion tool (GOrilla) to identify enrichment for gene ontology (GO) categories [167, 168].

### Screening for epistatic interactions

Epistatic hub genes were identified using MArginal ePIstasis Test (`MAPIT`), a linear mixed modeling approach that tests the significance of each SNP’s marginal effect on a chosen phenotype. `MAPIT` requires a complete genotype matrix, without missing data. SNPs were imputed using `BEAGLE 4.1` [169, 170] and then filtered for MAF ≥ 0.05 using `VCFtools` (v.0.1.16) [171]. `MAPIT` was run using the Davies method on the imputed genome (File S2), DGRP2 webserver-adjusted phenotype scores for each sex-based phenotype (S2 Table), DGRP2 relatedness matrix, and covariate file containing *Wolbachia* infection and the presence of five major inversions.

Resulting marginal effect *P*-values (data available File S3) were filtered to a Bonferroni threshold (*P* ≤ 2.56e-8) and tested for pairwise epistatic interactions in a set-by-all framework against the initial 1,901,174 SNPs (unimputed; MAF ≥ 0.05) using the `PLINK–epistasis` flag (v.1.90)[172]. Results were filtered for all *P*-values that exceeded a Bonferroni threshold, calculated as 0.05 / (the number of Bonferroni marginal effect *P*-values x 1,901,174 SNPs).

### Annotating FBgn and orthologs

Flybase gene (FBgn) identifiers were converted to their respective *D. melanogaster* (Dmel) or *H. sapiens* (Hsap) gene symbols using the *Drosophila* RNA^i^ Stock Center (DRSC) Integrative Ortholog Prediction Tool (DIOPT)[42]. FBgn were filtered for all high to moderate confidence genes, or low confidence genes if they contained the best forward and reverse score.

### Calculating an empirically simulated significance threshold

We sought to simulate an empirically derived significance threshold that was unique to our data set and separate from the traditional DGRP and Bonferroni thresholds used in other studies. Using the genotype-phenotype matrix, two separate datasets were simulated (n = 1000) for each sex-based phenotype. The first randomized the genotype-phenotype matrix using all available line means, while the second randomized subsets of 150 genotype-phenotype pairs.

Simulated associations were run with `PLINK` [172](v.1.90) on each dataset type for each sex-based phenotype. The 5^th^ percentile most-significant *P*-value across all permutations in a simulation type was deemed the “empirically simulated significance threshold.”

### GO term analysis

GOWINDA [49] was implemented to perform a Gene Ontology (GO) analysis that corrects for gene size in GWA studies. We conducted this analysis for male (n=418), female (n=473), sex-average (n=527), and sex-difference (n=214) candidate SNPs exceeding a relaxed P < 1E-4 significance threshold, against the 1,901,174 SNPs with MAF ≥ 0.05. We ran 100,000 simulations of GOWINDA using the gene mode and including all SNPs within 2000 bp.

Gene Ontology enRIchment anaLysis and visuaLizAtion tool (GOrilla)[167, 168] was run on `PEGASUS_flies` gene-scores and `Hierarchical Hotnet` sub-networks using the default commands and a gene list compiled from all genes available in the FB5.57 annotation file.

### Weighted Gene Co-expression Network Analysis

To test whether ambient adult transcriptomes could explain the observed phenotypic variation, we turned to the publically available DGRP2 microarray data, downloaded from the DGRP2 webserver [20]. These data represent the transcriptomes for untreated young adult flies of each sex. We performed Weighted Gene Co-expression Network Analysis (WGCNA) analyses using the available R package [63] to cluster and correlate the expression profiles of genes from 177 shared, DGRP lines. This analysis was run using the following parameters: power = 16 (from soft threshold analysis ≥ 0.9), merging threshold = 0.0, signed network type, maximum blocksize = 1000, minimum module size = 30.

### Data availability

All data required to rerun the outlined analyses either publically available through FlyBase (http://flybase.org/) [34, 35, 39], the DGRP2 webserver (http://dgrp2.gnets.ncsu.edu/), or in the Supplement: S1-S18 tables, https://doi.org/10.26300/v4rm-sa82; S1 file, https://doi.org/10.26300/dwvm-vt70; S2 file, https://doi.org/10.26300/317y-p682; S3 file, https://doi.org/10.26300/xcrh-c744; S4 file, https://doi.org/10.26300/qhc7-dp70.

## Acknowledgements

We thank Priya Nakka and Matthew A Reyna for their guidance through `PEGASUS` and `Hierarchical HotNet`, respectively. David Boerma for his assistance recording high-speed footage and F. Lemieux for maintaining *Drosophila* stocks. We are grateful for critical feedback from M. Tatar and T. Roberts and our anonymous reviewers.

## Author contributions

ANS, JAM, and DMR conceived and designed the study. ANS performed validation crosses, while ANS and JAM collected data. ANS performed the statistical analyses guidance from SR and LC. SPS and ANS designed and implemented `PEGASUS_flies`. ANS and DMR wrote the manuscript.

## Supplemental Results

### Establishing an empirically defined significance threshold

While the Bonferroni significance threshold is conservative, the conventional *P* = 1E-5 threshold might be considered lax. Accordingly, we simulated two sets of genotype-phenotype matrices; one set “shuffled” the genotype-phenotype matrix while the other set randomly “subsampled” 150 of the 197 lines.

The significance threshold for each sex-based phenotype in each simulation was determined by taking the 5^th^ percentile of the most significant *P*-value across 1000 permutations [173]. Despite these efforts, the resulting significance thresholds were even more stringent than the Bonferroni (S18 Table) and resulted in only one variant (2R_2718036_DEL) mapping to *CG15236* and *CG34215* in the shuffled sex-difference set. *CG15236*’s function is not well known, but it is expressed during embryogenesis and pupariation in the developing brain and central nervous system and putatively affects the wing veins [174, 175]. *CG34215* is less understood, though it is expressed at varying levels throughout developmental and adult stages [34] and contains a single domain Von Willebrand factor type C domain—thought to play a role in anti-viral capabilities [176].

## Supporting information

**S1 Fig. DGRP lines’ mean flight performance is highly repeatable across generations.** Set of genotypes (n = 12) reared 10 generations apart show very strong agreement (r = 0.95) in mean flight performance scores. The regression line (red line) through the point pairs (black points) has nearly the same slope and y-intercept as the x = y line (gray dashed line).

**S2 Fig. Sex-average and sex-difference phenotypic distributions are amenable to an association study.** Distribution in mean landing height (m) for (A) sex-average and (B) sex-difference phenotypes suggest ample phenotypic variation exists to run an association study. Each plot is sorted in order of increasing phenotype score, independent of one another.

**S3 Fig. QQ-plots show enrichment for some additive variants across each of the sex-based phenotypes.** Plots comparing the theoretical vs. observed *P*-value distribution across (A) males, (B) females, (C) sex-average, and (D) sex-difference phenotypes. Red line denotes y = x.

**S4 Fig. Top additive associations are spaced throughout the genome.** Top additive variants, those reported in DGRP2 webserver file with the ‘top.annot’ suffix, are largely free of linkage blocks. There is a larger block on X, corresponding with 10 variants that map to intronic and one synonymous coding site in *CG32506*. The heat component corresponds with likelihood of being in a linkage block from less (0 - blue) to more likely (1 - red).

**S5 Fig. Additional sex-based phenotype Manhattan plots for additive analysis.** (A) Males, (B) females, and (C) sex-difference phenotypes all have significant additive variants pass a traditional DGRP threshold (*P* ≤ 1E-5, gray solid line, red points), and at least one variant pass a Bonferroni threshold (*P* ≤ 2.63E-8, gray dashed line, red dot with black outline). Variants are arranged in order of relative genomic position by chromosome and plotted by the –log10 of the *P*-value. The sex-average is displayed in text.

**S6 Fig. Genetic crosses performed for deriving experimental and control stocks used to validate candidate genes.** All crosses are represented with females on the left and males on the right. Ten single pair crosses of a female genetic control, either w^1118^ (pictured) or y[1] w[67c23], in white boxes were crossed with the respective *Mi*{ET1} insertional mutant line in green boxes. After the initial cross, heterozygous flies were backcrossed to the respective genetic control for five generations. In the sixth generation, single pairs of heterozygous flies were crossed. Progeny without the Avic\GFP^E.3xP3^ marker were collected as homozygous nulls, while several vials of putatively homozygous mutants (no progeny without marker) were crossed again to confirm genotype. Stocks were monitored for two additional generations to confirm mutant carrier status before a homozygous mutant stock was selected as an experimental line.

**S7 Fig. Significant whole genes are distributed throughout the genome and sex-based phenotypes.** Whole gene analyses conducted with `PEGASUS_flies` for (A) males, (B) females, and (C) sex-difference phenotypes showed enrichment for significant whole genes across these three, and the sex-average (displayed in text). Each dot represents a whole gene, ordered by position across the chromosomes and plotted as the –log10 of the gene-score. Points above the Bonferroni threshold (*P* ≤ 3.03E-6, gray line) are colored in red.

**S8 Fig. Significant marginal variants are unevenly distributed across sex-based phenotypes.** (A) Males had very few significant variants pass a Bonferroni threshold (*P* ≤ 2.56E-8, gray solid line, red points), while (B) females had more and (C) sex-average had the most. (D) Sex-difference had no significant marginal variants. Variants are arranged in order of relative genomic position by chromosome and significance scores –log10 transformed.

**S9 Fig. Trait-relationship correlation matrix shows no correlation between measured phenotypes and young adult transcriptome**. Neither sexes’ mean landing height, standard deviation in landing height, or proportion of flies that fell through the column (fallen) were significant with a cluster of similarly expressed genes in a Weighted Gene Co-expression Network Analysis (WGCNA). Colored modules on the left represent WGCNA-generated clusters of genes and the color of each table cell corresponds with the magnitude of correlation coefficient (top number in cell). The bottom number in each cell is the significance of the correlation. No clusters were significantly correlated with any sex-phenotype combination.

**S1 Table.** *Drosophila* stocks used in this study.

**S2 Table.** Raw and adjusted flight performance metrics.

**S3 Table.** No significant correlations were observed between flight performance and other DGRP phenotypes.

**S4 Table.** Up to two inversions were significant covariates in three of the sex-based analyses.

**S5 Table.** Several additive variants associated with flight performance.

**S6 Table.** Several candidate genes were validated for flight performance.

**S7 Table.** Several gene-scores pass a Bonferroni threshold across all four sex-based phenotypes.

**S8 Table.** Twilight-estimated local False Discovery Rate (lFDR) cutoff thresholds for PEGASUS_flies gene-scores.

**S9 Table.** Hierarchical HotNet sub-network gene annotations.

**S10 Table.** Large sub-network from Hierarchical HotNet is enriched for trans-regulatory factors and neurodevelopmental Gene Ontology terms.

**S11 Table.** Collection of smaller sub-networks from Hierarchical HotNet are collectively enriched for mRNA splicing and autophagy Gene Ontology terms.

**S12 Table.** Significant marginal variants identified from MAginal ePIstasis Test (MAPIT).

**S13 Table.** Epistatic interactions play a large role in shaping the genetic architecture of flight performance.

**S14 Table.** Epistatic interactions with *pickpocket 23* (*ppk23*) are enriched in a Gene Ontology (GO) term analysis form neurodevelopmental genes.

**S15 Table.** Genes mapped to from epistatic interactions with *CG42671* are significantly enriched for neurodevelopment in a Gene Ontology (GO) analysis.

**S16 Table.** Gene set enrichment analysis for significant epistatic interactors within a 669 bp intergenic region between chr3L:6890373 - 6891042 suggests enrichment for neurodevelopmental Gene Ontology categories.

**S17 Table.** Master lookup table for all genes identified.

**S18 Table.** Empirically derived P-values from simulated permutations of the genotype-phenotype matrix.

**S1 File. High-speed gifs of strong, intermediate, and weak genotypes in flight performance assay**; https://doi.org/10.26300/dwvm-vt70

**S2 File. BEAGLE-imputed DGRP2 genome**; https://doi.org/10.26300/317y-p682

S3 File. All marginal *P*-value significance scores calculated using MAPIT; https://doi.org/10.26300/xcrh-c744

**S4 File. Source code for PEGASUS_flies**; https://doi.org/10.26300/qhc7-dp70 -- Also available at: https://github.com/ramachandran-lab/PEGASUS_flies

